# MRPL47 as a Novel Mitochondrial Biomarker for Early Detection and Therapeutic Response in Ovarian Cancer

**DOI:** 10.64898/2026.03.05.709884

**Authors:** Meenakshi Pradeep, Ishaque Pulikkal Kadamberi, Jasmine George, Yueying Gao, Camilla Dagum, Ajay Nair, Shirng-Wern Tsaih, Anjali Geethadevi, Anupama Nair, Elizabeth Hopp, Denise Uyar, William Bradley, Janet Rader, Yongsheng Li, Pradeep Chaluvally-Raghavan, Akinyemi I. Ojesina

## Abstract

MRPL47 (Mitochondrial Ribosomal Protein Large Subunit 47) gene in chromosome 3q26 encodes a protein that is part of the large subunit of the mitochondrial ribosome. We observed that MRPL47 is frequently amplified and overexpressed in ovarian cancer samples. Importantly, increased expression of *MRPL47* mRNA is associated with high levels of MRPL47 protein in ovarian cancer patients. High expression of MRPL47 is also associated with poor overall and recurrence free survival of ovarian cancer patients. Notably, MRPL47 improved metabolic fitness by enhancing cellular respiration, and glycolysis in cancer cells. Gene set enrichment analysis and target specific knockdown assays revealed that MYC transcription factor regulates MRPL47 expression. Furthermore, MRPL47 was identified very high in the plasma samples of ovarian cancer patients compared to those of healthy volunteers. MRPL47 was also associated with cisplatin resistance, whereas its expression predicted sensitivity to cisplatin therapy. Taken together, we demonstrated that MRPL47 can be used as a diagnostic biomarker for ovarian cancer and other cancers with 3q26 chromosomal amplification.

## Introduction

Ovarian cancer is challenging to detect early as it often remains asymptomatic until it reaches advanced stages and metastasizes. Therefore, improved screening methods are crucial for early detection and significantly enhancing patient survival rates ^1,2^. Currently, ovarian cancer diagnosis primarily relies on pelvic exam, various imaging modalities, and/or diagnostic laparoscopy for evaluating pelvic mass ^3,4^.

A commonly used and a well-accepted tumor biomarker for ovarian cancer is cancer antigen 125 (CA-125), often employed preoperatively to estimate the likelihood of malignancy ^5^. However, its effectiveness in detecting early-stage ovarian carcinoma is limited, as only 50% of stage I and II cases exhibit elevated CA-125 levels. Additionally, various benign ovarian conditions and non-gynecologic disorders can also cause increased CA-125 levels, reducing its specificity and sensitivity ^6,7^. These limitations suggest that CA-125 is a not a highly reliable biomarker for early detection and screening of ovarian cancer.

Our study aims to identify novel candidate proteins that can be useful for screening tests which can be executed with high-level accuracy with high sensitivity and specificity. Gene copy number alterations (CNAs), such as gene amplification are among the most frequent genomic alterations in ovarian cancer ^1^. Notably, amplification of the 3q26.3 locus occurs in approximately 30% of high-grade serous ovarian cancers (HGSOC) ^8–12^. While many genes in the 3q26 locus have been characterized for their mechanism and effects, the potential role of these genes and/or their encoded proteins as diagnostic biomarkers as well as their role in predicting the response to chemotherapy or immunotherapy has not been well established.

We hypothesize that certain low-molecular-weight proteins below 30 kDa are easily secreted into the circulation and therefore represent promising candidates for ovarian cancer diagnosis and for predicting the response against chemotherapy. In this study, we identified a low molecular weight protein MRPL47 (Mitochondrial Ribosomal Protein Large subunit 47) with the potential of developing as a biomarker for ovarian cancer detection and also for predicting the response for chemotherapeutics. MRPL47 is a protein-coding gene that is part of the large subunit of the mitochondrial ribosome (mitoribosome) ^13^. The mitochondrial ribosome is responsible for protein synthesis within mitochondria, which are organelles that produce adenosine triphosphate (ATP) and convert into energy in eukaryotic cells. The mitoribosome of mammals are 55-60S particles and are composed of small 28S and large 39S subunits ^13^.

Unlike other organelles, mitochondria contain their own genome, known as mitochondrial DNA (mtDNA), which encodes only 13 proteins. In contrast, all other mitochondrial ribosomal proteins (MRP’s) are encoded by nuclear genes. In humans, MRP gene family comprises of 30 genes which code for mitochondrial ribosomal small subunits (MRPS) and 50 genes for the large subunits (MRPLs) ^14^. Despite their essential role in mitochondrial function, MRPLs and MRPSs are poorly studied for their extra-mitochondrial functions and potential as a diagnostic or prognostic biomarker for cancer. Therefore, this study aims to investigate the role of MRPL47 as a potential diagnostic biomarker for ovarian cancer.

## Results

### MRPL47 copy number variations (CNV) are associated with high expression of MRPL47 in cancer

To identify the genes within the 3q26 locus that contribute to cancer oncogenesis, we evaluated the copy number variation profiles of lung squamous cell carcinoma (LUSC), cervical squamous cell carcinoma and endocervical adenocarcinoma (CESC), high grade serous ovarian carcinoma (HGSOC), lung adenocarcinoma (LUAD) (CESC), and uterine corpus endometrial carcinoma (UCEC), where CNVs are a frequent event in the TCGA dataset. Importantly, our analysis identified that MRPL47 is highly amplified, or copy-number gained as part of 3q26 locus in approximately or more than 50% patients of these cancer types, especially in HGSOC (**Fig. 1A and 1B**). Next, we determined if the CNV is a determinant factor of MRPL47 expression in those cancers where MRPL47 is highly amplified. Notably, our analysis identified that MRPL47 was significantly overexpressed in patients with copy number gain or amplification in all above cancers (LUSC expression data is not included due to space limit) (**Fig. 1C**). In addition, MRPL47 protein level was observed to be markedly higher in ovarian tumor samples compared to normal tissue samples in the clinical proteomic tumor analysis consortium (CPTAC) dataset (**Fig. 1D**). Next, we investigated the expression of MRPL47 at the protein level in a large number of clinical samples using tissue microarray (TMA) comprised of 118 pathologic samples including both ovarian cancer samples and normal tissue samples. Consistent with the CPTAC data, our TMA staining revealed that MRPL47 was highly expressed in ovarian cancer samples compared to both normal and benign tissue samples (**Fig. 1E, 1F, 1G and Supplementary Fig-1**).

**Figure 1.**
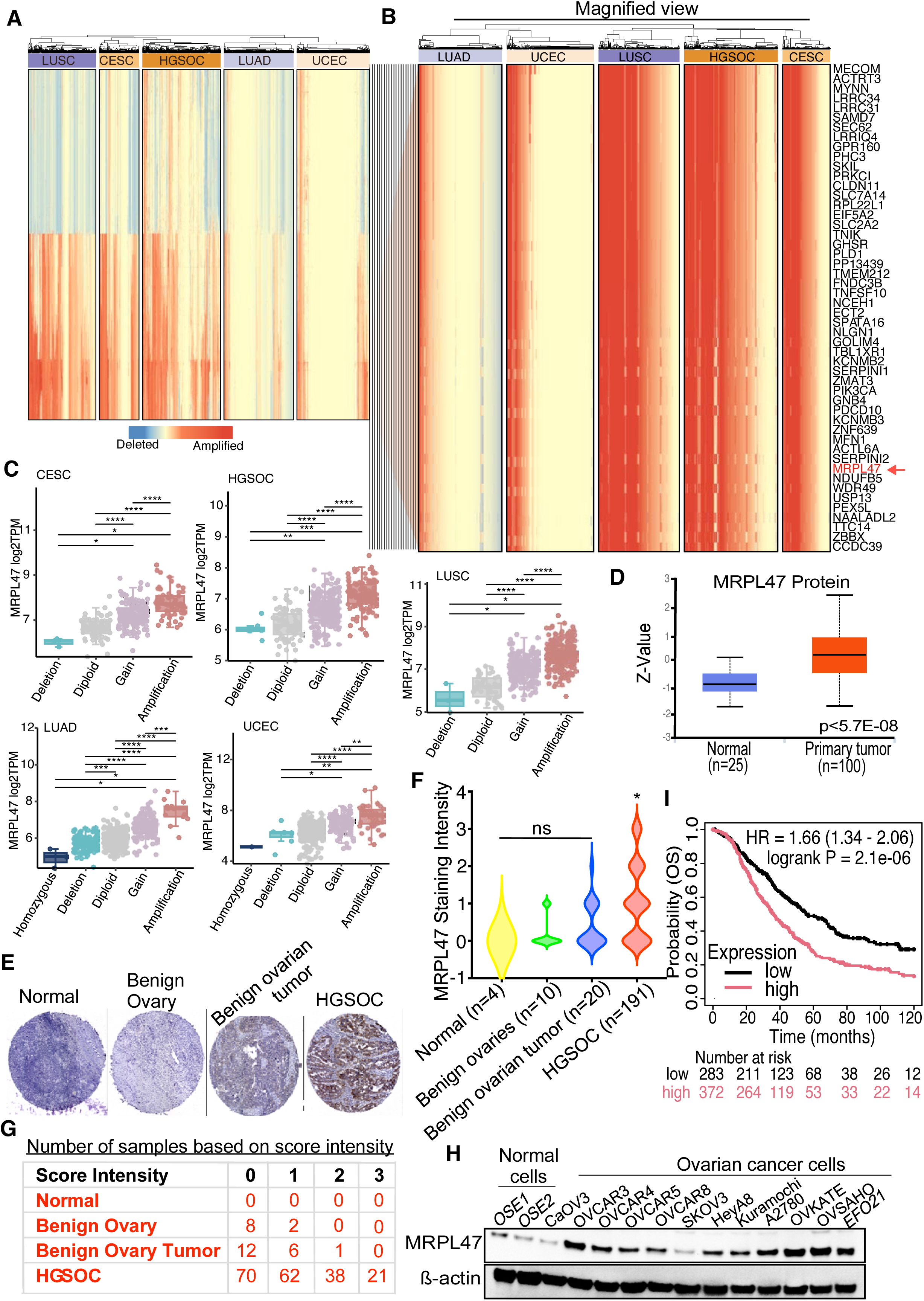
Copy number variation (CNV) is associated with MRPL47 expression. (**A**) The copy number profiles of CESC (Cervical Squamous Cell Carcinoma and Endocervical Adenocarcinoma, n=295), LUAD (Lung Adenocarcinoma, n=516), LUSC (Lung Squamous Cell Carcinoma, n=501), HGSOC (High Grade Serous Ovarian Serous Ovarian Carcinoma, n=579) and UCEC (Uterine Corpus Endometrial Carcinoma, n=539) in chromosome 3. Red represents copy number amplification, and blue represents copy number deletion. (**B**) The copy number profiles of top 50 genes with the most copy number amplification. (**C**) The distribution of MRPL47 expression across diverse groups with different copy number levels of MRPL47, including homozygous deletion (Homozygous), single copy deletion (Deletion), diploid normal copy (Diploid), low-level copy number amplification (Gain) or high-level copy number amplification (Amplification). Comparisons between groups were performed via a two-tailed Wilcoxon rank sum test. The levels of significance are denoted as *p < 0.05, **p < 0.01, ***p < 0.001, ****p < 0.0001. **D,** Box plots show the level of MRPL47 protein in the ovarian cancer patients and normal ovary in the publicly available dataset of Clinical Proteomic Tumor Analysis Consortium. **E,** Representative images of MRPL47 immunohistochemistry was performed on normal, benign ovary (ovaries diagnosed with non-cancer diseases), benign tumors and HGSOC tissue samples (n=225) on two tissue microarray (TMA) slides. **F,** MRPL47 staining intensity score in all sample types. The levels of significance are denoted as *p < 0.05, ns, non-significant. **G,** Table shows the tumor intensity score in each sample group **H**, Indicated cell lines including ovarian surface epithelial (OSE) cells and ovarian cancer cells were lysed and immunoblotted for MRPL47 and beta-Actin. **I**, Overall survival (OS) based on MRPL47 mRNA expression in ovarian cancer patient samples was determined. Samples were stratified into high and low MRPL47 mRNA expression groups. Significance between groups were determined by using Log Rank statistical test.

Immunoblot analysis of lysates from normal human ovarian surface epithelial (OSE) cells and ovarian cancer cells further showed that most of the HGSOC cell lines had markedly elevated levels of MRPL47, whereas CaOV3 and SKOV3 expressed very modest levels of MRPL47. As expected, the normal OSEs expressed very low levels of MRPL47 (**Fig. 1H**). Next, we sought to determine whether MRPL47 mRNA levels were associated with patient outcomes in a publicly available ovarian cancer dataset by stratifying patients into low and high expression group. We found that high MRPL47 mRNA expression was associated with worse overall survival (OS) in the patient group whose tumors express higher levels of MRPL47 (**Fig. 1I**).

### MRPL47 expression is regulated by transcription factors in ovarian cancer

Next, we investigated which genes and cancer hallmark pathways are related to the MRPL47 gene expression in ovarian cancer. For this, we performed the gene set enrichment analysis (GSEA) analysis ^15^ by ranking all the protein-coding genes based on their gene expression correlation with MRPL47 expression, and against the cancer hallmark pathways gene set. Importantly, our analysis identified that most of the genes positively correlated with MRPL47 in the TCGA dataset were classified under the annotation of MYC target genes, oxidative phosphorylation, MTORC signaling and glycolysis (**Fig. 2A).** This data prompted us to evaluate if MYC has any important role in regulating MRPL47 expression. As expected, silencing of MYC reduced the levels of MRPL47 in OVCAR8 and MDAMB-231 cancer cell lines (**Fig. 2B**). We also performed an immunofluorescence staining for MYC and MRPL47 in ovarian cancer tissues and found that MRPL47 expression was higher in the cells that express high levels of MYC, which again suggested that MYC transcription factor could regulate the transcription of MRPL47 (**Fig. 2C**). As a complementary approach to knocking down MYC, we also used a MYC inhibitor (Z,E)-5-(4-Ethylbenzylidine)-2-thioxothiazolidin-4-one (a.k.a. 10058-F4) and found that the treatment with MYC inhibitor consistently reduced the levels of MRPL47 in both OVCAR8 and MDAMB231 cells (**Fig. 2D**). The transcription factor binding site determining program JASPAR identified MYC consensus sequence (**Fig. 2E**) in MRPL47 promoter sites (**Supplementary Fig 2A**). Importantly, our chromatin immunoprecipitation (ChIP) assay followed by qPCR also showed enrichment of MYC on MRPL47 promoter markedly (**Fig. 2F, 2G**). Taken together, our data show that MYC transcription factor is an important regulator of MRPL47 in ovarian cancer cells (**Fig. 2H**).

**Figure 2.**
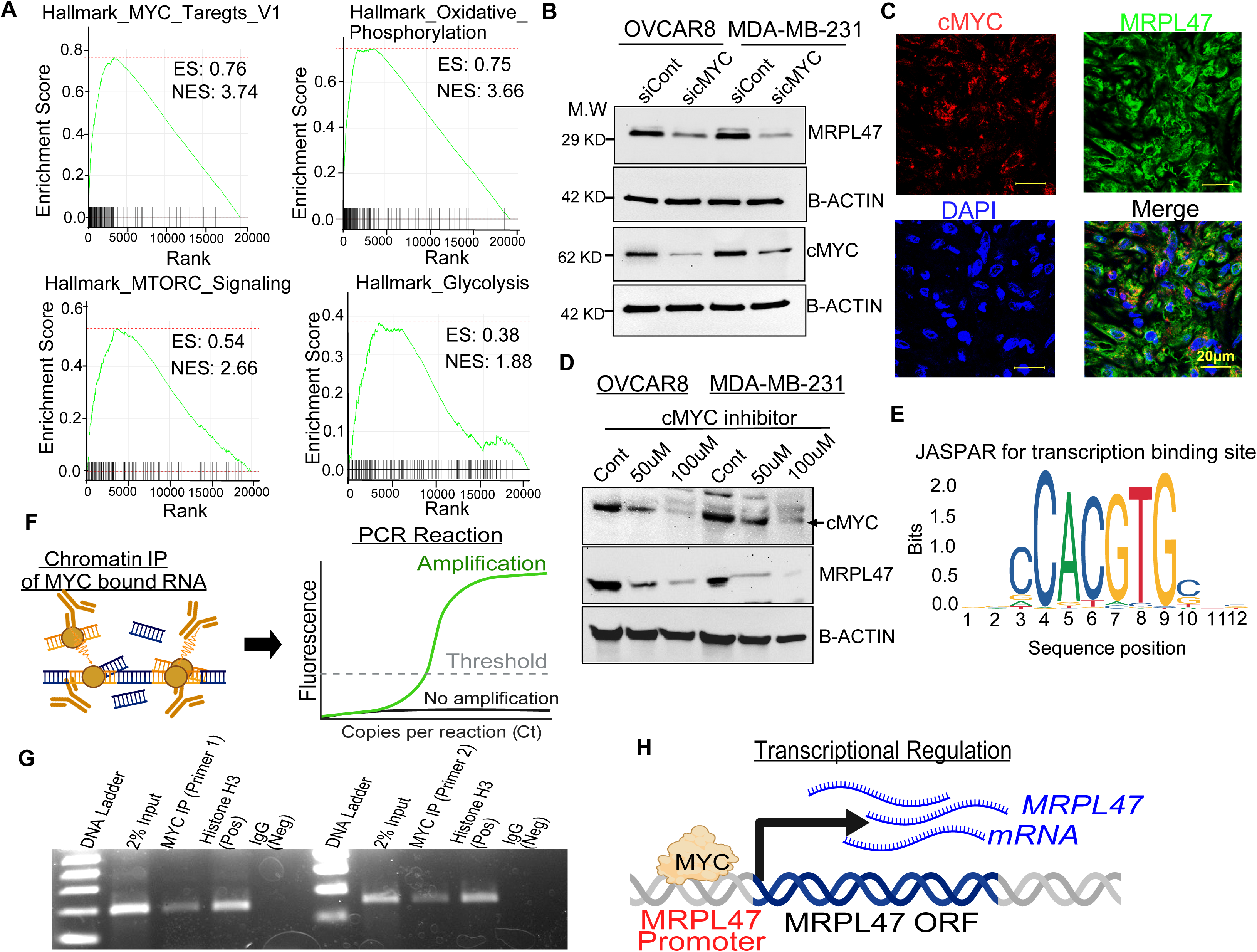
MRPL47 is regulated by transcription factor cMYC. **A,** Gene Set Enrichment Analysis demonstrates the enrichment score (ES) of indicated cancer hallmarks based on the genes are correlated with MRPL47 expression in the TCGA ovarian cancer samples. ES: enrichment score. **B,** Ovarian cancer cell line OVCAR8 and breast cancer cell line MDA-MB-231 were transfected with pre-validated MYC (cMYC) siRNA (5nM) for 48hours, then Western blot was performed to detect MRPL47 and MYC using target specific antibodies. β-Actin was used as an internal control. **C,** Immunocytochemistry was performed in ovarian tumor tissues using MRPL47 Anti-rabbit and cMYC (Anti-mouse) antibodies. **D,** OVCAR8 and MDA-MB-231 cells were treated with MYC inhibitor 10058-F4 for 24h then lysed and immunoblotted using indicated antibodies. **E,** Consensus sequence of the cMYC predicted by JASPAR database **, F,** Schema shows how PCR was performed after chromatin-IP of cMYC. **G,** qPCR products of elutes of cMYC chromatin IP were then resolved in 2% agarose gel and photographed. **H,** Model shows how MRPL47 is transcriptionally regulated by MYC.

### MRPL47 is highly expressed in patient plasma samples and elevated MRPL47 levels are associated with cisplatin resistance

Based on our data that MRPL47 is highly amplified and expressed in most of the ovarian cancer patients (**Fig.1**), we hypothesized that MRPL47 could be used as a diagnostic biomarker for ovarian cancer. To explore this, we aimed to determine the levels of MRPL47 in the plasma from ovarian cancer patients and compare them to those in the plasma of healthy individuals (**Table 1**). Blood samples were collected from patients and healthy controls, then plasma were separated and enzyme linked sorbent assay (ELISA) was performed. Notably, our ELISA analysis using 15 samples each from both healthy controls and ovarian cancer patients identified that MRPL47 is highly expressed in ovarian cancer patient samples compared to the healthy controls (**Fig. 3A**).

**Figure 3.**
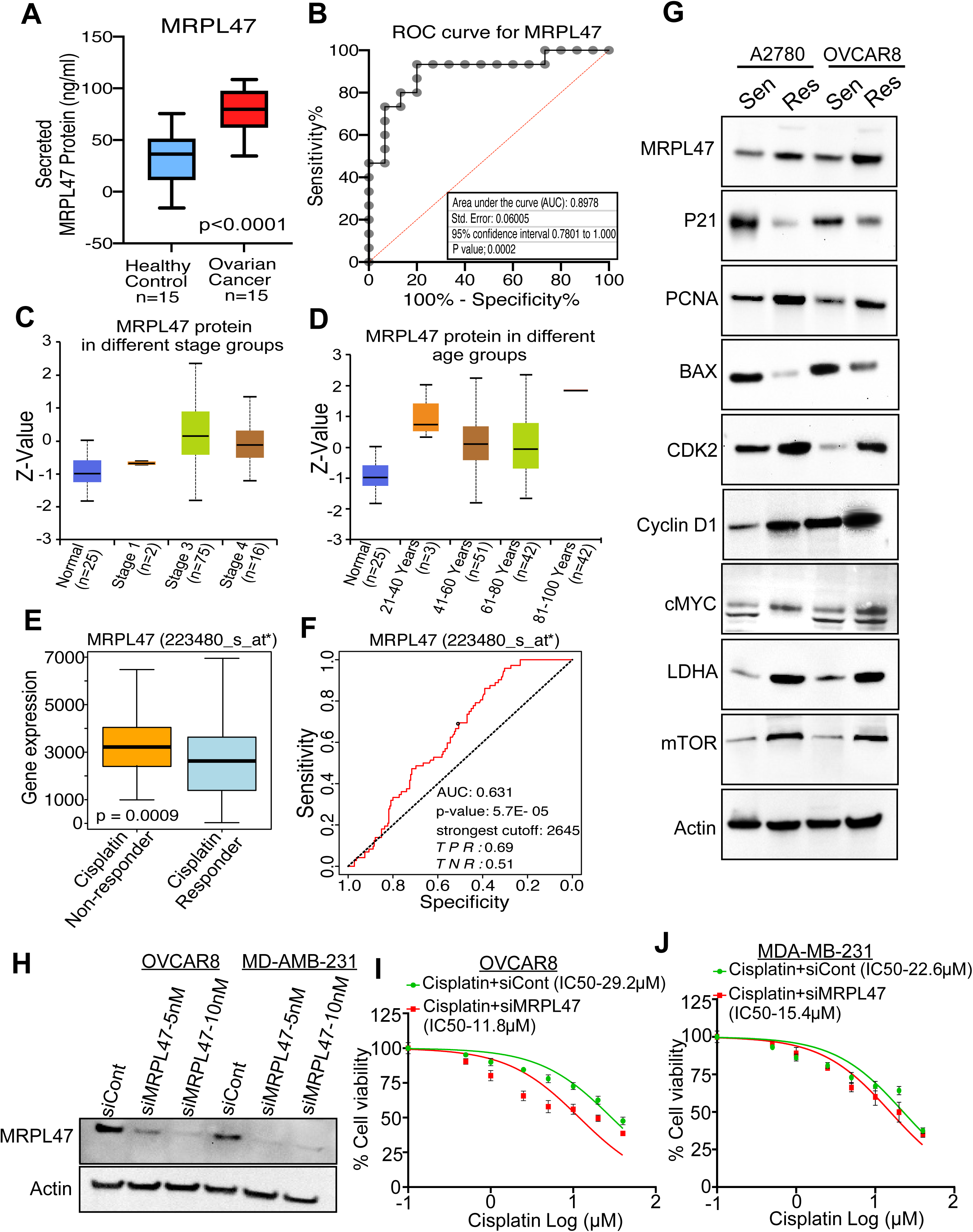
MRPL47 is significantly elevated in the plasma samples of ovarian cancer patients and MRPL47 level is associated with chemoresistance. **A,** ELISA quantification of MRPL47 levels in plasma from healthy controls and ovarian cancer patient samples. **B**, Receiving operating characteristics (ROC) curve analysis for MRPL47 detection using ELISA. AUC, area under the curve. **C-D**, Box plots shows MRPL47 protein levels in different stages and age group of ovarian cancer patients. **E-F**, The ROC analysis based on MRPL47 expression and platin therapy determined using publicly available online ROC analysis plotter ^31^. **G,** Parent and cisplatin resistant versions of A2780 and OVCAR8 cells were lysed and immunoblotted using indicated antibodies. **H,** Ovarian cancer cell line OVCAR8 and breast cancer cell line MDA-MB-231 were transfected with indicated concentrations of pre-validated MRPL47 siRNA for 48hours, then Western blot was performed using indicated antibodies. B-Actin was used as an internal control. **I-J,** OVCAR8 and MDAMB231 cells were either transfected with control siRNA (green line) or MRPL47 siRNA (red line) for 24h and then treated with cisplatin up to 40µM concentration and cell viability was determined using CCK8 assay.

**Table 1.** Characteristics of Patient Population.

| **Table 1** Characteristics of Patient Population |  |  |
| --- | --- | --- |
| Characteristics | Normal (n) | HGSOC(n) |
| Age (y) |  |  |
| ≤40 | 13 | 0 |
| 41-50 | 2 | 0 |
| 51-60 | 0 | 3 |
| ≥60 | 0 | 12 |
| Race |  |  |
| White | 10 | 14 |
| Black | 0 | 0 |
| Hispanic | 0 | 0 |
| Other | 5 | 1 |
| Patient Stage | None | Stage-3B* |

Next, we constructed a logistic regression model and the ROC curve to determine if the relative fold change in the levels of MRPL47 could effectively differentiate between ovarian cancer patients and healthy controls (**Fig. 3B**). The area under the curve (AUC) for MRPL47 was 0.8978, with a 95% confidence interval (CI) ranging from 0.78 to 1.0, indicating high sensitivity and specificity in distinguishing between the two groups (**Fig. 3B**). These findings demonstrate that MRPL47 can serve as a reliable diagnostic biomarker for ovarian cancer, offering high sensitivity and specificity for distinguishing cancer patients from healthy controls. We also found that MRPL47 expression is higher in the patients who are in late stages of ovarian cancer (**Fig. 3C**). Strikingly, we found that the patients in the younger age group between 21 years to 40 years expressed higher level of MRPL47 compared to the patients in either 41-60 years or 61-80 years (**Fig. 3D**). The need for novel proteins to identify chemotherapy resistance like platinum-based therapy is critical because many ovarian cancer treatments fail due to resistance to platinum agents in young age patients. Therefore, relying on known biomarkers is often insufficient, as cancer cells employ diverse and complex mechanisms to survive upon chemotherapy. Thus, we sought to determine if MRPL47 expression was associated with chemotherapy responses. Importantly, our expression analysis and corresponding ROC analysis identified that MRPL47 is very high in the samples collected from the patients who were not responding to platinum-based therapy (**Fig. 3E**).

The corresponding AUC analysis of the therapy response dataset yielded a value of 0.631 with a true positive rate (TPR) of 0.69 and true negative rate (TNR) of 0.51 respectively (**Fig. 3F**) again confirming the link between MRPL47 expression and platinum-based therapy sensitivity. To further corroborate the role of MRPL47 expression in cell line models, we examined the expression of MRPL47, and proteins associated with cell survival and apoptosis in parent and platinum resistant ovarian cancer cells. Importantly we found cisplatin resistant A2780 and OVCAR8 cell lines expressed high levels of MRPL47, and high levels of proteins associated with cell survival such as PCNA, CDK2, LDHA, mTOR and Cyclin D1, and low levels of proteins associated with cell cycle control or apoptosis such as Bax, and p21 (**Fig. 3G**). Complementary to the observation that MRPL47 is associated with resistance to chemotherapeutics, knockdown of MRPL47 using target-specific siRNA indeed improved sensitivity to cisplatin (**Fig. 3H-3J**). Herein, we observed that the IC50 of cisplatin is reduced approximately 60% in OVCAR8 cells and approximately 30% in highly aggressive, metastatic and difficult to treat MDAMB231 cell line (**Fig. 3H-3J**).

### MRPL47 promotes a hybrid metabolic phenotype with high level glycolysis and oxygen consumption rate in tumor cells

Because MRPL47 is a mitochondrial protein with the potential to reshape mitochondria-driven cancer cell metabolism, we sought to overexpress MRPL47 in two cell lines SKOV3 and MDA-MB-468, which express low levels of MRPL47, and metabolic flux was analyzed using the Seahorse XFe96 bioenergetic analyzer. Notably, MRPL47 overexpression resulted into a more metabolically active cells than the control group. We found that MRPL47 increased the glycolytic capacity in MDA-MB-468 and SKOV3 cancer cells (**Fig. 4A**). Further analysis revealed that mitochondrial basal OCR, maximal OCR, and ATP turnover were all significantly elevated when MRPL47 is overexpressed (**Fig. 4B**). Likewise, extracellular acidification rate (ECAR) measured after oligomycin treatment reflected that glycolytic capacity was significantly higher in MRPL47-overexpressing SKOV3 cells and increased slightly in MDA-MB-468_MRPL47 cells, indicating enhanced glucose catabolism independent of mitochondrial oxygen consumption (**Fig. 4C**). The glycolytic reserve, defined as the difference between basal ECAR and ECAR following oligomycin addition, was also significantly increased in both cell lines, suggesting a metabolic shift toward a more energetic and glycolytic state (**Fig. 4D**). Collectively, our results suggest that MRPL47 is important for cancer cell metabolism, which makes the cancer cells more metabolically active by allowing cells to utilize both glycolysis and oxidative phosphorylation to meet high energy demands for cellular proliferation, and chemoresistance.

**Figure 4.**
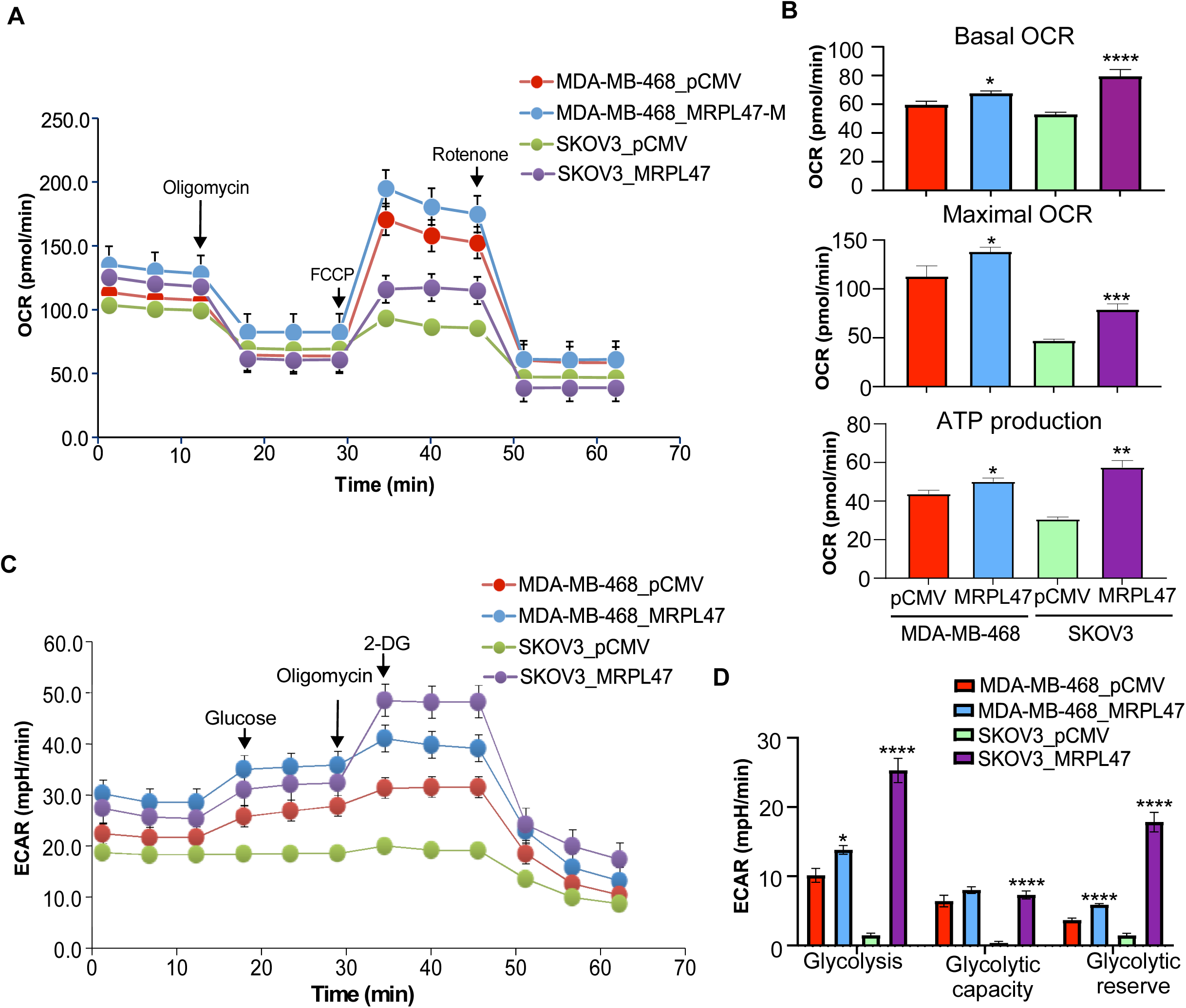
Overexpression of MRPL47 leads to hybrid metabolic phenotype with high glycolysis and improved oxygen consumption rate. **A,** Oxygen consumption rate (OCR) in MDA-MB-468 and SKOV3 cell lines were transfected with MRPL47 expressing plasmid (pCMV3-MRPL47-OFPSpark) and its vector control pCMV3 plasmid using the Seahorse XFe96 extracellular flux analyzer. **B,** Bar graph shows the results of mitochondrial respiration changes analyzed with basal respiration, maximal respiration, and ATP production. **C,** Extracellular acidification rate (ECAR) measurement in the cells from A **D,** Bar graph shows indicated glycolytic functions in the cells were transfected with MRPL47 expressing plasmid (pCMV3-MRPL47-OFPSpark) and its vector control pCMV3 plasmid. Mean ± SEM (n at least 10 from biological replicates). *p < 0.05, **p < 0.01, ***p < 0.000, ****p < 0.0001, ns: non-significant.

## Discussion

The amplification of the 3q26 locus is a common genetic alteration found in various cancers, particularly those with aggressive phenotypes and poor prognoses ^11,16^. This amplification results in the overexpression of oncogenes and regulatory elements that drive tumorigenesis, metastasis, and therapy resistance in cancers such as ovarian cancer, lung squamous cell carcinoma, head and neck squamous cell carcinoma, cervical cancer, and endometrial cancer. Among these, 3q26 amplification is most prevalent in ovarian cancer. Key oncogenes, including PIK3CA (Phosphatidylinositol-4,5-Bisphosphate 3-Kinase, Catalytic Subunit Alpha), TERC (Telomerase RNA Component), PRKC1 (Protein Kinase C iota), and SOX2 (Sex-Determining Region Y-Box 2), are located within the 3q26 locus ^10,16–18^. Additionally, MRPL47, which resides in this locus, shows genomic amplification and is associated with elevated mRNA and protein expression in ovarian cancer tissue samples.

Mitochondria contain a distinct family of proteins called MRPs, which are essential components of the 55S mitochondrial ribosome, made up of a small (28S) and a large (39S) subunit ^13,14^. MRPs are encoded by nuclear genes, and the proteins they produce are translocated to the mitochondria, where they help translate the 13 essential oxidative phosphorylation proteins encoded by mitochondrial DNA (mtDNA) ^14^. Therefore, these proteins are critical for the electron transport chain (ETC) and ATP production via oxidative phosphorylation. MRPL47 is a component of the 39S large subunit of the mitochondrial ribosome (mitoribosome).

Our study suggests that increased levels of MRPL47 due to gene copy amplification improves mitochondrial function in cancer cells, which in turn supports the survival and growth of tumor cells sporadically. Emerging evidence highlights that several MRPLs play critical roles in regulating cancer metabolism and tumor progression. Beyond their canonical functions in mitochondrial translation, our data shows that MRPLs can influence tumor cell metabolism to favor tumor growth with an increase in glycolysis, oxidative phosphorylation, and lipid biosynthesis, thereby energizing the tumor cells for hypergrowth and chemoresistance. Ovarian cancer cells, particularly HGSOC, exhibit increased mitochondrial respiration and ATP generation ^19^. Enhanced mitochondrial function plays a crucial role in ovarian cancer progression by fueling tumor metabolism, promoting survival under stress, and contributing to therapy resistance. Consistent with this notion, we demonstrate that MRPL47 overexpression enhances both glycolytic and mitochondrial respiratory activity, with a hybrid metabolic state that supports rapid cell growth.

It has also been reported that MYC protein in cancer cells with a hybrid glycolysis/OXPHOS metabolic phenotype enables metabolic flexibility, allowing cancer cells to utilize diverse nutrients including glucose, fatty acids, and glutamine to meet energy and biosynthetic demands across varying microenvironments ^20^. Notably, we found that MYC transcription controls the transcription of MRPL47, revealing a regulatory mechanism underlying the hybrid metabolic phenotype in cancer cells potentially operated through MYC and MRPL47.

While high ECAR alone is typical of the Warburg effect in most of the cancer cells, some cancers exhibit a hybrid metabolic state where both glycolysis and oxidative phosphorylation are active, and both are increased relative to normal cells. Likewise, MRPL47 overexpression exhibits high rate of both processes simultaneously, which can be indicative of a highly adaptable and metabolically flexible phenotype. Presumably, this hybrid metabolic state allows them to adapt to different nutrient availabilities and microenvironments, potentially contributing to increased chemoresistance. Similar oncogenic functions have been reported for other MRPL family members, such as MRPL13, MRPL15, and MRPL35, which have been shown to sustain mitochondrial integrity, elevate ATP production, and drive anabolic metabolism in diverse cancer types^21–24^.

While cytoplasmic ribosomal proteins of the 80S ribosome (60S and 40S subunits) have been extensively characterized for their roles in general protein synthesis, the potential of using those proteins as both diagnostic or prognostic marker for ovarian and other cancers are not well established. Importantly, the increased levels of MRPL47 in plasma samples demonstrated that MRPL47 can be used as a diagnostic biomarker for ovarian cancer. The tumors collected from the patients in later stages also exhibited high levels of MRPL47. Furthermore, high levels of MRPL47 in ovarian cancer patients in the younger age group underscore the significance of protein levels as biomarkers for young age cancer lies in their ability to aid in early detection, and its application to monitor treatment, and predict prognosis, even though this area of research is still developing in our laboratory.

Importantly, the ELISA data in conjunction with AUC analysis suggests that MRPL47 may be investigated as a diagnostic biomarker for identifying patients who are less likely to respond to standard platinum-based chemotherapy. This may allow for early stratification of patients and enable the design of personalized treatment strategies. Moreover, the strong correlation between MRPL47 upregulation and enhanced expression of pro-survival and metabolic regulators and the finding that MRPL47 silencing restores cisplatin sensitivity underscores its therapeutic relevance, suggesting that targeting MRPL47 could improve chemotherapy efficacy and overcome drug resistance in ovarian and possibly other aggressive cancers. Collectively, these results position MRPL47 as a promising biomarker on a panel that also includes CA 125 and actionable molecular target with both diagnostic and therapeutic value in clinical oncology. Our analysis have also shown that MRPL47 is highly amplified and overexpressed in several other malignancies including lung, head and neck, cervical, and endometrial cancers (**Fig-1**) suggesting that MRPL47 could be developed as a diagnostic biomarker and also as a biomarker that can predict the outcome of chemotherapy in those cancers too.

## METHODS

### Genomic Data Analysis

Data from The Cancer Genome Atlas (TCGA) were first analyzed using cBioPortal (http://www.cbioportal.org/). The segmented data for all the samples were downloaded from the Broad GDAC Firehose portal (http://firebrowse.org, version: 20160128), followed by standard GISTIC2 (Genomic Identification of Significant Targets in Cancer, version 2) analysis using Firehose-suggested parameters.

### Patient Samples

All clinical samples used in this study were collected in deidentified manner according to the Institutional Review Board approved protocol PRO00011631 at Medical College of Wisconsin after obtaining informed consent. Normal ovarian tissues were sourced from the tumor-free ovary of patients with unilateral ovarian cancer, patients with other gynecological cancers not involving the ovary. Similarly, blood samples were collected in deidentified manner for plasma preparation. Control blood samples were obtained from healthy individuals with no history of gynecological diseases. All study subjects were U.S. residents and provided written informed consent before sample collection, in accordance with the Institutional Review Board- approved protocol at the Medical College of Wisconsin.

### Cell Culture

Kuramochi cells were received from Taru Muranen at Beth Israel Deaconess Medical Center, Boston, MA, USA. NIH-OVCAR8 cells were purchased from the National Cancer Institute (NCI) cell line repository. SKOV3, MDA-MB-231, and MDA-MB-468 cells were purchased from ATCC/PBCF repository. All cancer cell lines were cultured in RPMI-1640 medium (ATCC, Manasassas, VA, USA) supplemented with 10% fetal bovine serum (FBS, Omega Scientific), 1% Antibiotics (Anti-Anti 100X, Thermo Scientific). Cells were routinely tested and deemed mycoplasma free of using the Plasmotest^TM^ Mycoplasma Detection Kit (InvivoGen, San Diego, CA). Authenticity of the cell lines used were confirmed by STR characterization at IDEXX Bioanalytics Services (Columbia, MO).

### Tissue Microarray Immunohistochemistry

MRPL47 protein levels in human ovarian cancer tissues were analyzed using two TMAs slides. First, slides were dewaxed in xylene and rehydrated through graded ethanol to distilled water. Antigen retrieval for the slide specimens were performed using IHC-Tek epitope retrieval solution and steamer set (IHC World, LLC.). The slides were then immersed in 3% H_2_O_2_ for 10 min to quench endogenous peroxidase followed by blocking with 10% goat serum for 1 h. Slides were then stained overnight at 4°C with the MRPL47 primary antibody (#PA5-65479, Thermo Fisher Scientific). Following DAB staining (Vector Labs, Burlingame, CA), the slides were counterstained with Harris modified hematoxylin (Thermo Fisher Scientific Inc., Rockford, IL), dehydrated with graded ethanol and xylene, and finally mounted with paramount. TMAs slides were digitally scanned using Pannoramic 250 FLASH III scanner (3D HISTECH ltd. Version 2.0) and, using the Case Viewer software (3D HISTECH ltd. Version 2.0) was used to view and analyze images.

### Gene Set Enrichment Analysis (GSEA)

The Cancer Genome Atlas (TCGA) ovarian cancer gene expression data were analyzed using cBioPortal ^25^. We selected the ‘Ovarian Serous Cystadenocarcinoma (TCGA, PanCancer Atlas)’ dataset, then queried for MRPL47 gene, chose ‘Co-expression’ tab, and selected batch corrected RSEM normalized mRNA expression data to obtain the Spearman’s rank correlation of all the protein coding genes to MRPL47. This list of protein coding genes ranked in the ascending order of their correlation coefficient was used as input for the GSEA analysis using fgsea R package ^26^. The enrichment score (ES) was calculated for each functional set, reflecting the extent to which a gene set is overrepresented at the top or bottom of the ranked list. The normalized enrichment score (NES) was calculated based on 1000 permutations. Cancer hallmark gene sets from MSigDB and the gene sets with false discovery rate <0.001 were considered as selection criteria. The cancer hallmark pathways from MSigDB ^27^ was used as the gene set input to fgsea. We used the following parameters for the fgsea analysis: p value boundary parameter ‘eps’ = 0; minimal gene set size ‘minSize’ = 10; and maximal gene set size ‘maxSize’ = 500.

### siRNA Transfection

All siRNA duplexes for human cMYC, MRPL47 and scrambled control were purchased from Ambion. Reverse transfections were performed using the Lipofectamine RNAiMAX transfection reagent (Thermo Fisher Scientific Inc., Waltham, MA). At 48 hrs post-transfection, cells were harvested for further analysis.

### MRPL47 overexpression

To perform overexpression of MRPL47 in SKOV3 and MDA-MB-231 cells, we transfected the cells with control pCMV3 vector or MRPL47 expressing pCMV3-MRPL47-OFPSpark (Sino Biological) using Lipofectamine 2000 (Invitrogen, Carlsbad, CA). Cells were collected 48h after transfection and Western blotting was performed to check the expression of MRPL47 and other targets in cells followed by Seahorse assay.

### Western Blot Analysis

Western blot analysis was performed using precast gradient gels (Bio-Rad, Hercules, CA) following standard methods. Briefly, cultured cells were lysed in RIPA buffer containing protease inhibitors and phosphatase inhibitors (Santa Cruz Biotechnology, Dallas, TX). Protein concentration was determined by Pierce BCA Protein Assay kit (Thermo Fisher Scientific, Waltham, MA). 25 μg of protein was separated by SDS-PAGE and transferred onto a PVDF membrane (Bio-Rad, Hercules, CA). The membranes were then probed with the primary antibodies, followed by peroxidase-conjugated secondary antibody. Chemiluminescent signals were developed using BioRad’s Chemiluminescence imager.

### Cell Counting Kit-8 (CCK-8) assay

Cell viability was assessed using the CCK-8 assay (Dojindo Molecular Technologies, Inc.), according to the manufacturer’s protocol. Briefly, OVCAR8 and MDA-MB-231 cells treated with siCont and MRPL47 siRNA (5nM) for 24 h were re-seeded (10x103 cells/well) into 96-well plates. Following dose-dependent treatment with cisplatin (0-40µM), 10 μl CCK-8 solution was added to each well and incubated for 2 h at room temperature. The absorbance value of each well at a wavelength of 450 nm was measured using an microtiter plate reader (Tecan, Mannedorf, Switzerland).

### Enzyme Linked Immunosorbent Assay (ELISA)

ELISA tests were performed to quantify MRPL47 levels in plasma samples from healthy individuals and ovarian cancer patients. The Human 39S Ribosomal Protein L47 (MRPL47) ELISA Kit was purchased from Abbexa Ltd (Catalog No: abx542868). Plasma samples were collected from individuals after informed consent according to an Institutional Review Board approved protocol at Medical College of Wisconsin. Samples were centrifuged at 1000 × g for 20 minutes to remove precipitate and then diluted with diluent at 1:10 before ELISA. 100 μl of diluted samples were then added into the wells and mixed using a microplate shaker. ELISA plates were incubated at 37°C for 2 hours using a microplate shaker. Following incubation, the samples were discarded, and the wells were treated sequentially with the primary antibody containing Detection Reagent A, followed by the HRP-conjugated secondary antibody with Detection Reagent B, as per the manufacturer’s instructions. After thorough washing, Tetramethylbenzidine (TMB) substrate solution was added, followed by a stop solution. Optical density (OD) was measured immediately at 450 nm.

### Chromatin Immunoprecipitation (CHIP) Assay

The ChIP assay was performed as outlined previously ^28,29^ with some modifications using SimpleChIP® Enzymatic Chromatin IP Kit (Magnetic Beads) (Cell Signaling Technologies, #9003). In brief, cells were seeded in 150 mm dish and crosslinked by adding 37% formaldehyde to a final concentration of 1% for 15 min at room temperature followed by quenching with glycine. Subsequently, cells were washed with PBS and subjected to cell lysis using cell lysis buffer provided in the kit. The cell extracts were sonicated using Bioruptor at 4 °C and DNA was extracted and ran on 2% agarose gel to check the appropriate shearing. The lysates were immunoprecipitated with cMYC antibody (Cell Signaling Technologies, #18583) and DNA was isolated and purified. DNA obtained was quantitated and qPCR was performed using the designed MRPL47 promoter region primers (**Supplementary Table S1**).

### Measurement of Oxygen Consumption Ratio (OCR)

Mitochondrial OCR was measured in an XFe96 Extracellular Flux Analyzer (Agilent Technologies, Santa Clara, CA, USA) as previously shown ^30^ in MRPL47 overexpressed MDA-MB468 and SKOV3 cells. In brief, cells transfected with control and MRPL47 plasmids were seeded in XF96 cell culture microplate (Agilent Technologies). Prior to the experiment, the culture media was replaced with Agilent Seahorse XF RPMI Medium supplemented with D-Glucose Pyruvate and glutamine. Three basal OCR measurements were recorded prior to injection of oligomycin. After recording the oligomycin-sensitive OCR, FCCP-sensitive rates were recorded. Finally, antimycin A/rotenone was injected to inhibit electron flow through the electron transport system. The data were normalized relative to the protein concentration in each well and are presented as the percentage of change as compared with control.

### Measurement of Extracellular Acidification Rate (ECAR)

The ECAR was measured in a similar way in an XFe96 Extracellular Flux Analyzer in both MRPL47 overexpressed MDA-MB-468 and SKOV3 cells. ECAR measurements were recorded followed by an injection of a saturating level of glucose, measuring a glucose-induced response. This was followed by Oligomycin A, an ATP synthase inhibitor, which inhibits mitochondrial ATP production and shifts the energy production to glycolysis, revealing the cellular maximum glycolytic capacity. Finally, 2-deoxyglucose, a glucose analog, was injected to inhibit glucose binding to hexokinase, thus confirming extracellular rates to be a product of glycolysis. Basal glycolysis and glycolytic capacity were calculated and normalized relative to the concentration of protein in each of the corresponding microplate wells. All the data were compared to and expressed as percent of control.

### Statistical Analysis

Data are presented as means ± Standard Error (SEM) or Standard Deviation (SD). Statistical comparisons were performed using unpaired two-tailed Student’s t-tests or Wilcoxon rank sum test, where appropriate, with a probability value of 0.05 considered significant. Cox proportional hazard regression model was used for univariate survival analysis. Either overall survival or recurrence-free survival was used as an endpoint. The cutoffs of high expression and low expression were optimized to achieve the lowest p value. Statistical analysis was performed using R Software or GraphPad Prism Software. Logistic regression analysis was used to develop the model for diagnosing cancer and predicting treatment outcomes, and the receiver operating characteristic (ROC) curve was employed to evaluate its performance.

## Author contributions

Conceptualization, Writing—Original Draft and Figure Preparation, Investigation, M.P., J.G.; I.P.K., Bioinformatics Analysis, Y.G., Y.L., A.I.O, S-W, T., Ajay. N., P.C.-R., Tissue Microarray Development and clinical Data Analysis, C.D., D.U., W.B., E.H., J.R., Writing—Review & Editing, M.P., J.G.; Ajay. N., I.P.K., A.G., Anupama, N., D.U., J.R., Y.L., A.I.O, P.C.-R. Funding Acquisition, and Supervision, A.I.O, P.C.-R.

## Legends to Supplementary Figures

**Supplementary Figure-1.** Photographs of the two slides of TMAs of human ovarian cancer tissues used for immunohistochemical staining for MRPL47. The slides were stained for MRPL47 and scanned using an Aperio Scan Scope (Aperio Technologies).

**Supplementary Figure 2. A,** Human MRPL47 promoter sequence highlighted with MYC binding sites predicted by JASPER. **B,** Chromatin fragments prepared after sonication DNA fragments were resolved in an Agarose gel (1.5%) to determine the fragmentation of sonication.

**Supplementary Table S1.**
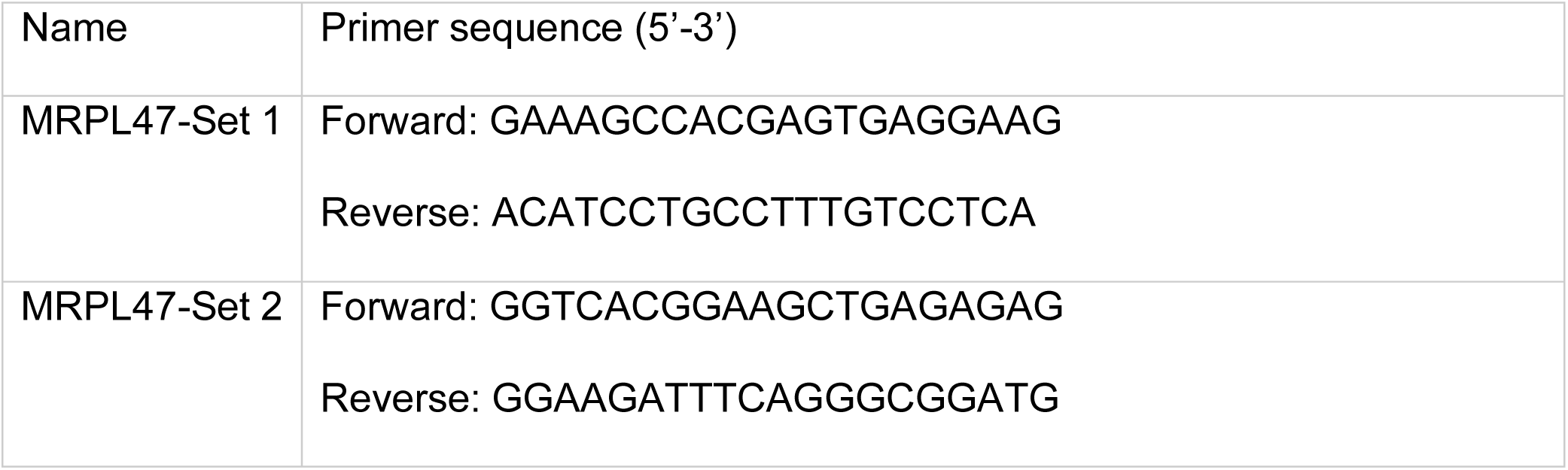
List of qRT-PCR promoter primer sequences.

