## Supplementary figures and images for "MRPL47 as a Novel Mitochondrial Biomarker for Early Detection and Therapeutic Response in Ovarian Cancer"

### Supplementary Figure-1

Supplementary Figure 1

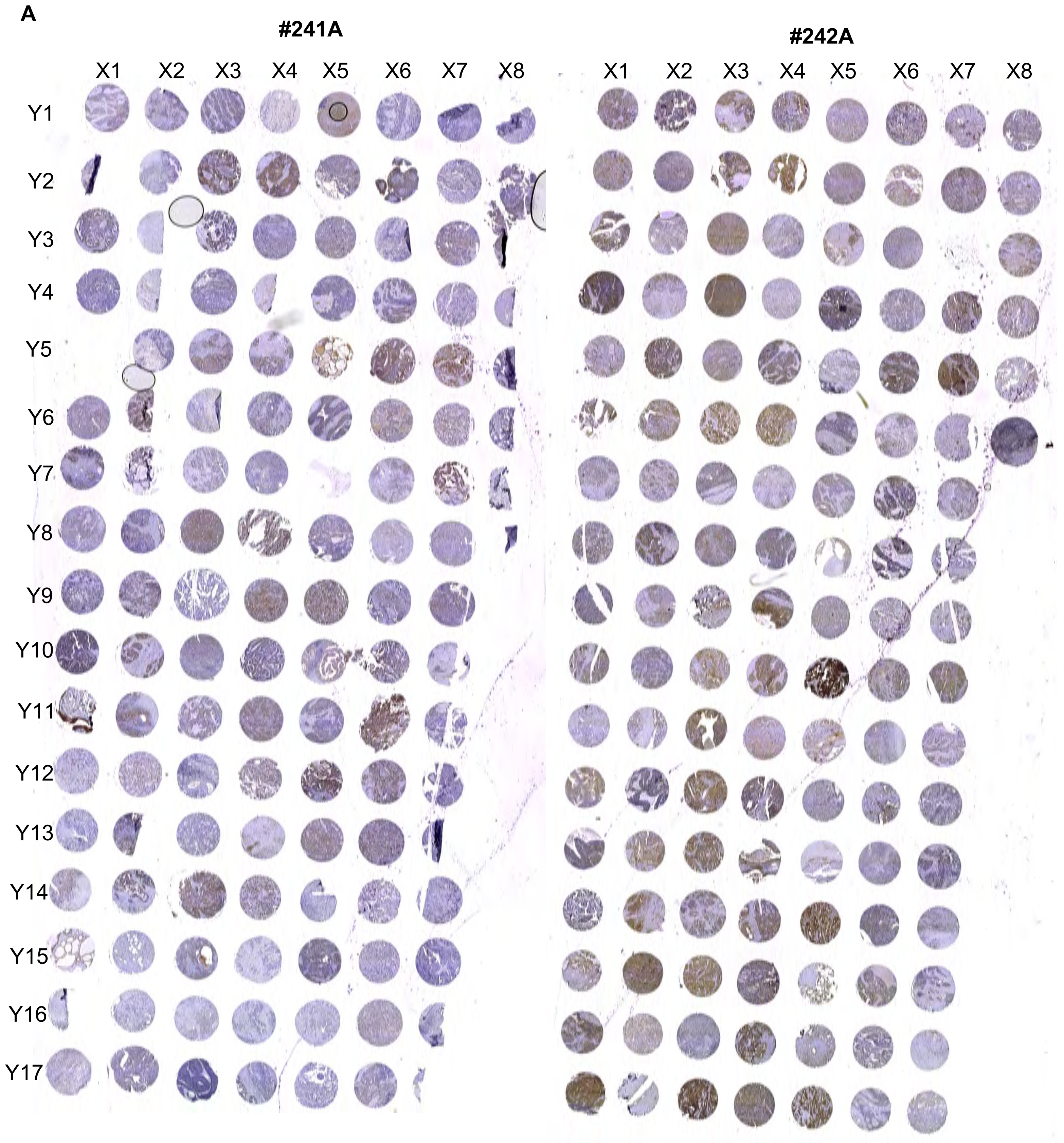
