## Supplementary Figure-2 for "MRPL47 as a Novel Mitochondrial Biomarker for Early Detection and Therapeutic Response in Ovarian Cancer"

**A**

**MRPL47 mitochondrial ribosomal protein L47 [ Homo sapiens (human)] Promoter Sequnce**

**<u>lower case indicates  sequences of upstream of Transcription Start Site</u>**

gctaataatgtaaacacgatctatctggctacttcaggaaaacccctcatatagctcgtg
ctccgtctctcctgtcaagccccaagcccacaacttgtcccggctccctccggatcccac
caaggtcaccctggaagcgtcctcccttttctcgccccgcacccgcttcacgcccgttct
ccctaggccagcacccaccagcagccttcggaaagccacgagtgaggaagccccccaaatc
cgaggcgagtgccaaggggccggccagacagagctgccaccgcagtaaccgaaacccgcc
gcaacaaactcatggccgccatggctactacgggcaggaggaagaagggaggcgggtcac
ggaagctgagagagccaatttaccaatgaggacaaaggcaggatgtggccatcttaagtg
agggcaagttttctattctcctcaccgccggcttgagacgccggaagtgacgctatcgtt
ggcggaaacgcgttttgccAGTTATGCGAAAACATGGCTGCGGCCGGTTTGGCCCTTCTT
TGTAGGAGAGTTTCATCCGCCCTGAAATCTTCCCGATCGTTAATAACTCCTCAGGTCCCT

**B**

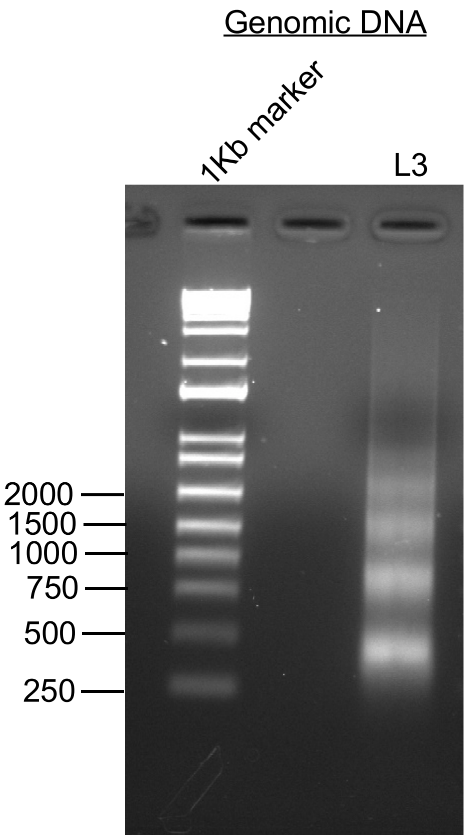
